# Kun-peng: an ultra-memory-efficient, fast, and accurate pan-domain taxonomic classifier for all

**DOI:** 10.1101/2024.12.19.629356

**Authors:** Qiong Chen, Boliang Zhang, Chen Peng, Jiajun Huang, Xiaotao Shen, Chao Jiang

## Abstract

Comprehensive metagenomic sequence classification of diverse environmental samples faces significant computing memory challenges due to exponentially expanding genome databases. Here, we present Kun-peng, featuring a unique ordered 4GB block database design for ultra-efficient resource management, faster processing, and higher accuracy. When benchmarked on mock communities (Amos HiLo, Mixed, and NIST) against Kraken2, Centrifuge, and Sylph. Kun-peng matched Sylph, achieving the highest precision and lowest false-positive rates while demonstrating superior time and memory efficiency among all tested tools. Furthermore, Kun-peng’s efficient database architecture enables the practical utilization of large-scale reference databases that were previously computationally prohibitive. In comprehensive testing across 586 air, water, soil, and human metagenomic samples using an expansive pan-domain database (204,477 genomes, 4.3TB), Kun-peng classified 69.78-94.29% of reads, achieving 38-43% higher classification rates than Kraken2 with the standard database. Unexpectedly, Sylph failed to classify any reads in air samples and left > 99.85% of reads unclassified in water and soil samples. In terms of computational efficiency, Kun-peng processed each sample in 0.2∼11.2 minutes using only 4.0∼35.4GB peak memory. Remarkably, these processing times were comparable to Kraken2 using the standard database (81GB, 5% of the pan-domain database). Memory-wise, Kun-peng required only 35.4GB peak memory with the pan-domain database, representing a 473-fold reduction compared to Kraken2. When compared to Sylph, Kun-peng processes samples up to 46.3 times faster while using up to 20.6 times less memory. Overall, Kun-peng offers an ultra-memory-efficient, fast, and accurate solution for pan-domain metagenomic classifications.

## Main

Metagenomics, a revolutionary genomic tool that bypasses traditional culturing methods, has transformed our understanding of microbial worlds by studying collective genetic material from mixed communities of organisms. This approach offers unprecedented insights into microbial diversity, ecological niches, genetic and evolutionary relationships, and functional activities, with applications spanning environmental science, bioremediation, bioenergy, and human health ^1–3^. By facilitating the understanding of soil microorganisms’ biogeochemical processes, metagenomics enables better climate impact predictions and strategies to combat warming and soil degradation^4,5^. Metagenomics is essential for the One Health approach, deepening our comprehension of the intricate relationships between microbes, humans, animals, and the environment. As it evolves, metagenomics continues to catalyze innovations across disciplines, offering integrated solutions that promote the health of all living systems on our planet. However, comprehensive analysis of metagenomic data, especially from non-human sources, faces significant challenges due to a vast number of uncultured and unknown species.

Moreover, metagenomic sequencing has also been used to evaluate the biological diversity of animals, plants, and other single-celled eukaryotes such as fungi ^6,7^. A useful strategy for these complex samples would be expanding the reference database to include as many isolated and metagenome-assembled genomes (MAGs) as possible. Still, the computational memory required to handle such large databases, which can easily surpass terabytes, is becoming increasingly unwieldy for most mainstream computing platforms.

Existing metagenomic analysis tools present various trade-offs in addressing these challenges^8,9^. These tools can be broadly categorized into database-based and machine-learning methods. Database-based approaches primarily include alignment-based, marker-based, and k-mer based methods ^10^. For example, alignment-based methods, such as BLAST, offer high sensitivity but are very computationally expensive ^11^. Marker-based methods, exemplified by MetaPhlAn 4, are fast and accurate with low memory usage but are limited to bacteria, archaea, and selected single-celled eukaryotes ^12^. K-mer-based methods, such as Kraken2, excel in speed with a relatively low memory footprint but struggle with large reference databases and increased false positive rates ^13^. Researchers have proposed improvements to address the limitations of k-mer-based methods, including opting for specific data structures and statistical approaches. For instance, Centrifuge’s FM index for efficient k-mer searches ^8,14^ and MegaBlast’s seed search strategy ^15^, but these methods introduce new trade-offs between speed and accuracy. The recently published Sylph tool introduces k-mer subsampling (default 1/200) with zero-inflated Poisson statistics for metagenome profiling. While this method achieves high precision on simple prokaryotic benchmark datasets and reduces computational overhead ^16^, it shows dramatically reduced sensitivity when profiling complex environmental samples in this study. Machine-learning techniques, including IDTAXA ^17^ and DeepVirFinder ^18^, show promise but generally underperform compared to database-based methods in classification accuracy ^10^. In theory, database-based methods can achieve higher classification sensitivity and accuracy when supported by curated, comprehensive reference databases ^10^. Nevertheless, the primary challenge for database-based tools remains their substantial memory requirements, particularly for complex environmental microbiome and exposome studies necessitating massive reference databases ^7,19^. For example, even the standard Kraken2 database requires approximately 100 GB of memory to run, surpassing the capabilities of most affordable personal computers or workstations, limiting its accessibility in institutions without high-performance computing platforms (HPCP). A highly memory-efficient tool is required to democratize metagenomics across the global scientific community.

Here, we introduce Kun-peng (https://github.com/eric9n/Kun-peng), an ultra-memory-efficient metagenomic classification tool (Fig. 1). Inspired by Kraken2’s k-mer-based approach, Kun-peng employs algorithms for minimizer generation, hash table querying, and classification. The cornerstone of Kun-peng’s memory efficiency lies in its unique ordered block design for reference database. This strategy dramatically reduces memory usage and improves processing speed, enabling Kun-peng to be executed on both personal computers and HPCP for most databases. Moreover, Kun-peng incorporates an advanced sliding window algorithm for reference database construction and sequence classifications to reduce false-positive rates. Finally, Kun-peng supports parallel processing algorithms to further bolster its speed. Kun-peng offers two classification modes: Memory-Efficient Mode (Kun-peng-M) and Full-Speed Mode (Kun-peng-F). Remarkably, Kun-peng-M processes classification faster than Kraken2 while requiring 11.56 to 473-fold less memory, with memory savings scaling proportionally to reference database size. Kun-peng-F loads all the database blocks simultaneously, matching Kraken2’s memory usage while achieving even greater speed improvements than Kun-peng-M, processing data in less than half the time required by Kraken2. Notably, Kun-peng is compatible with the reference database built by Kraken2 and the associated abundance estimate tool Bracken ^20^, making the transition from Kraken2 effortless. The name “Kun-peng” was derived from Chinese mythology and refers to a creature transforming between a giant fish (Kun) and a giant bird (Peng), reflecting the software’s flexibility in navigating complex metagenomic data landscapes.

**Fig. 1.**
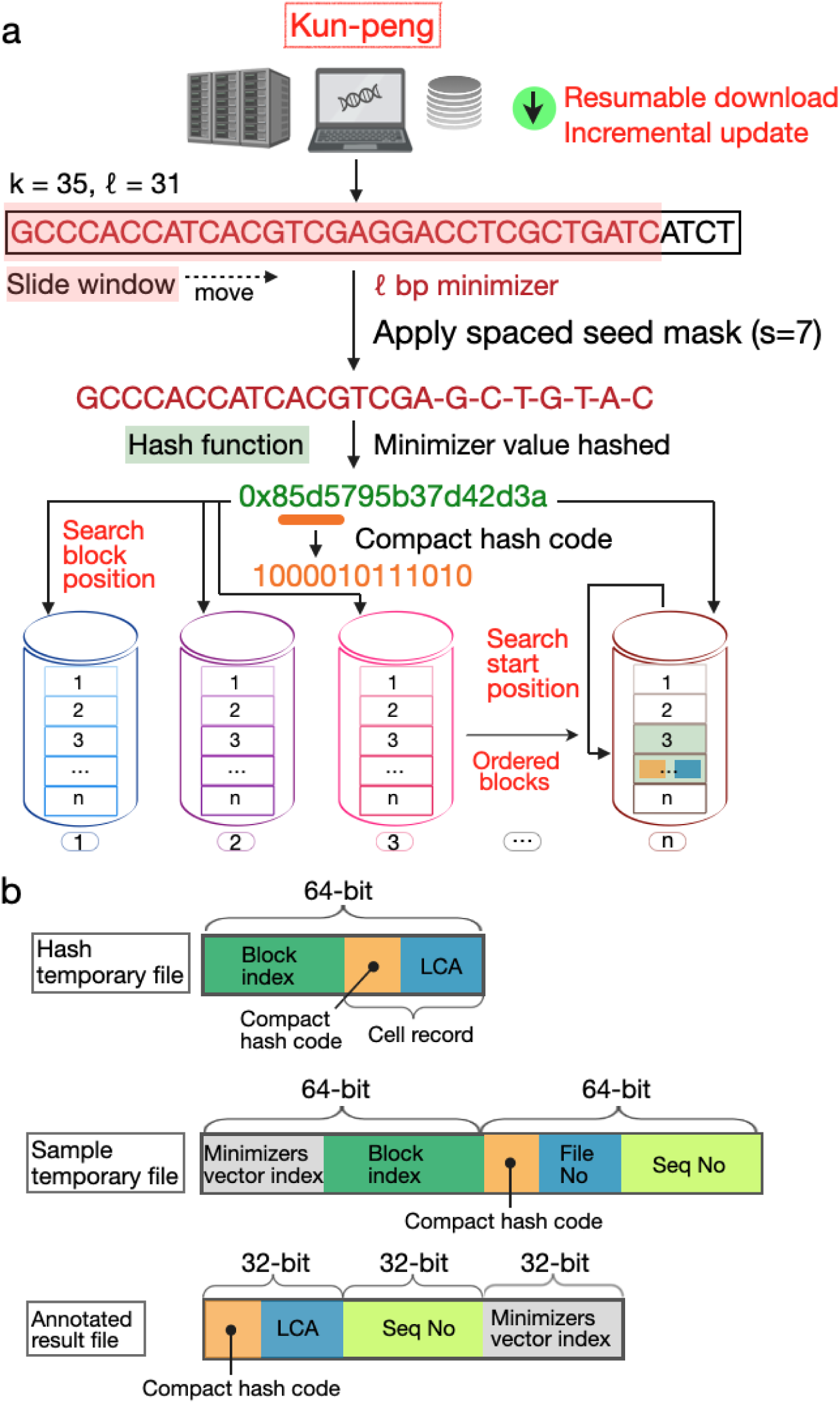
Overview of Kun-peng’s data retrieval, minimizer generation, and block-based database architecture. a. Kun-peng’s workflow from data acquisition to database searching. The pipeline begins with resumable download and incremental update capabilities for retrieving genomic data from databases. For sequence processing, a k=35 bp k-mer is analyzed using a sliding window to extract an ℓ=31 bp minimizer (red box). Unlike Kraken2, Kun-peng retains only unique ℓ bp minimizers to prevent over-counting and reduce false positives. A spaced seed mask (s=7) is applied to the minimizer sequence, followed by hash function computation to generate a compact hash code. This hash code determines both the block position for storage and the search start position within ordered blocks in the database. The database is divided into multiple ordered blocks (1 to n), enabling efficient memory usage through block-wise loading and searching. b. Data structure design of three key file types in Kun-peng: Hash temporary file: 64-bit structure containing block index, compact hash code, and LCA (Lowest Common Ancestor) information; Sample temporary file: 64-bit×2 structure storing minimizer vector index, block index, compact hash code, file number, and sequence number; Annotated result file: 32-bit×3 structure comprising compact hash code, LCA, sequence number, and minimizer vector index.

To assess Kun-peng’s accuracy and performance, we used two datasets comprising 14 mock metagenomes ^21,22^, varying in GC content (42.92% ∼ 53.94%), size (228M ∼ 474M in.gz formats), and composition characteristics (Table S1 and S2). We benchmarked Kun-peng against three tools that support custom databases and operate with relatively low memory requirements: Kraken2, Centrifuge, and Sylph. MegaBLAST was tested but abandoned due to excessive time requirements (>10 days per dataset). We did not compare Kun-peng to Metaphlan4 and GTDB because these two methods heavily relied on marker gene sets and cannot be easily extended to viruses, animals, and plants. All tools were run with default parameters except for Kraken2 and Kun-peng, where the confidence score was set to 0.3. Taxonomic classifications and abundance estimates for Kraken2, Kun-peng, and Centrifuge were processed with Bracken ^20^. As most classifiers considered below 0.01% abundance false positives, we removed these taxa for simplicity ^8^. Notably, Kun-peng exhibited identical performance utilizing a partitioned or unpartitioned database (Fig. S1).

Critical metrics for metagenomic classification include precision, sensitivity, and the area under the precision-recall curve (AUPRC). After filtering the low-abundance species, these metrics were calculated at the genus or species level. All tools demonstrated superior performance at the genus level compared to the species level (Fig. 2a). At both taxonomic levels, Kun-peng, Kraken2, and Sylph achieved comparable precision, while Sylph exhibited higher sensitivity than both Kun-peng and Kraken2. Centrifuge consistently showed the lowest precision among all tested tools (Figure 2a). We focused on the genus level due to higher overall performance. Kun-peng and Sylph demonstrated equally low false-positive rates, while Kraken2 showed higher false positives compared to Sylph. Statistical analysis revealed no significant difference in false-positive rates between Kun-peng and Sylph (Fig. 2b). Centrifuge’s high false positives likely result from lossy database compression (Fig. 2b). Kraken2’s higher false positives stem from redundant ℓ bp minimizer counts in the same read positions during classification (Fig. 2b; Methods). To further evaluate performance, we calculated the accuracy for estimating relative abundance for all samples. Kun-peng achieved comparable accuracy to Kraken2 in abundance estimation (Fig. 2c)

**Fig. 2.**
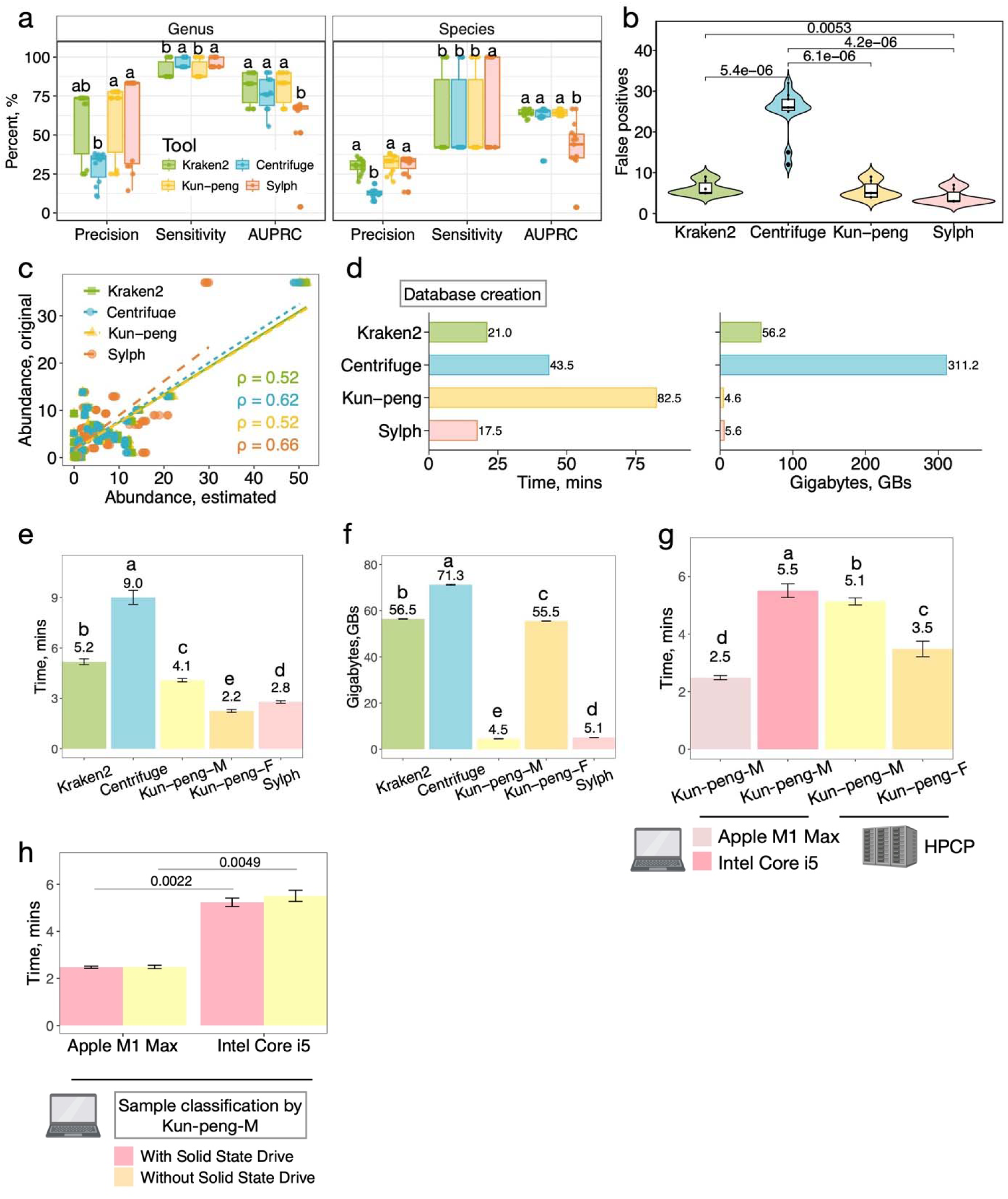
Comprehensive performance evaluation of Kun-peng against state-of-the-art metagenomic classifiers on mock communities. a. Classification performance at genus and species levels showing precision, sensitivity, and AUPRC (Area Under the Precision-Recall Curve) scores. Different letters indicate significant differences (p < 0.05) based on the Kruskal-Wallis test with post hoc Games-Howell tests. b. False positive rates comparison at genus level. P-values from pairwise comparisons using the Kruskal-Wallis test with post hoc Games-Howell tests are shown. c. Correlation between estimated and true genus-level abundances, with Spearman correlation coefficients (ρ) shown for each tool. d. Computational requirements for database construction (42,485 genomes), showing processing time (left) and peak memory usage (right) for each tool. e-f. Performance comparison for processing BMock12 community samples ^23^ (six paired-end samples, averaging 25 million reads): peak memory usage I and processing time (f). Different letters indicate significant differences (p < 0.05). g. Processing time comparison of Kun-peng modes (Memory-Efficient: Kun-peng-M; Full-Speed: Kun-peng-F) across different computing platforms (Apple M1 Max, Intel Core i5, and HPCP). Different letters indicate significant differences (p < 0.05). h. Impact of solid-state drive (SSD) on Kun-peng-M processing time across different computing platforms. P-values from the Kruskal-Wallis test with post hoc Games-Howell tests are shown.

We constructed a standard database using the complete RefSeq genomes of archaeal, bacterial, and viral domains. The database contains 123GB of fasta sequences, generating different index sizes across tools: 56GB for Kraken2 and Kun-peng, 72GB for Centrifuge, and 4.7GB for Sylph. Importantly, Kun-peng partitions the 56GB hash index into fourteen 4GB sub-hash blocks. Database construction time for Kun-peng was noticeably longer due to sub-hashing calculations (Fig. 2d). However, Kun-peng’s peak memory usage during database construction was only 4.6GB, representing just 8.19% and 1.50% of the memory required by Kraken2 and Centrifuge, respectively. Notably, this was even lower than Sylph’s memory usage (5.6GB) despite Sylph’s aggressive 1/200 k-mer subsampling approach (Fig. 2d).

Kun-peng operates in two modes for taxonomic classification: Memory-Efficient Mode (Kun-peng-M) and Full-Speed Mode (Kun-peng-F), both delivering identical classification results. In Memory-Efficient Mode, Kun-peng-M demonstrates remarkable resource efficiency, requiring only 4.5 ± 1.1 GB peak memory - representing just 7.96 ± 0.19%, 6.31 ± 0.15%, and 88.7 ± 2.06% of the memory used by Kraken2, Centrifuge, and Sylph, respectively (Fig. 2e). Despite this minimal memory footprint, Kun-peng-M maintains competitive processing speeds, using only 78.9 ± 3.67% of Kraken2’s and 45.3 ± 1.80% of Centrifuge’s processing time (Fig. 2f).

The Full-Speed Mode Kun-peng-F was optimized for processing speed while maintaining reasonable memory usage. Compared to Kraken2, it reduces processing time to 42.3 ± 2.47% while matching memory usage. Compared to Centrifuge, Kun-peng-F uses 77.9 ± 0.22% of the memory while requiring only 24.9 ± 1.16% of the processing time (Fig. 2e, f). Notably, Kun-peng-F achieves 80.78% ± 3.55% faster processing speed than Sylph (Fig. 2e, f).

Remarkably, with an ultra-low memory requirement, Kun-peng-M can even operate on most personal computers when the standard reference database is used (Fig. 2g). The storage limitations for samples and database can be addressed by equipping personal computers with an external hard drive without impacting processing time (Fig. 2h).

Environmental metagenomic data pose significant challenges for taxonomy classifications. Expansive pan-domain databases enhance classification coverage in environmental microbiome and exposome studies. However, the substantial memory requirements of current tools often impede efficient taxonomical classifications with expansive reference databases. Kun-peng addresses this challenge with its unique format of reference database. To demonstrate its capabilities, we constructed an expansive pan-domain database encompassing over 204,477 species (4.3TB of fna sequences), which generated a 1.85TB reference database for Kun-peng (Kun-peng-epdb) and an 81GB reference database for Sylph (Sylph-epdb). In contrast, both Kraken2 and Centrifuge failed to construct their extended pan-domain databases due to excessive memory requirements exceeding 1.85 TB. Standard databases provided by Kraken2 (Kraken2-sdb, 81GB) and Sylph (Sylph-sdb, 14GB) were also included in the comparisons. Using this expansive pan-domain database, Kun-peng-epdb achieved robust classification rates (69.78∼94.29%) and surpassed Kraken2-sdb by 43.4% ± 18.2%, 41.0% ± 21.5%, 38.8% ± 9.15%, and 7.40% ± 5.94% for samples from air, water, soil, and human body sites, respectively, while enhancing the overall diversity of classified taxa (Fig. 3a-3h). Notably, Sylph-sdb showed severely limited classification capability, leaving >90% of reads unclassified across all sample types (99.96% air, 91.60% water, 95.15% soil, 91.60% human). Surprisingly, Sylph-epdb performed even worse - failing to classify any reads in all 301 air samples and leaving out 99.98%, 99.85%, and 80.25% of reads unclassified in water (n=105), soil (n=135), and human (n=45) samples, respectively.

**Fig. 3.**
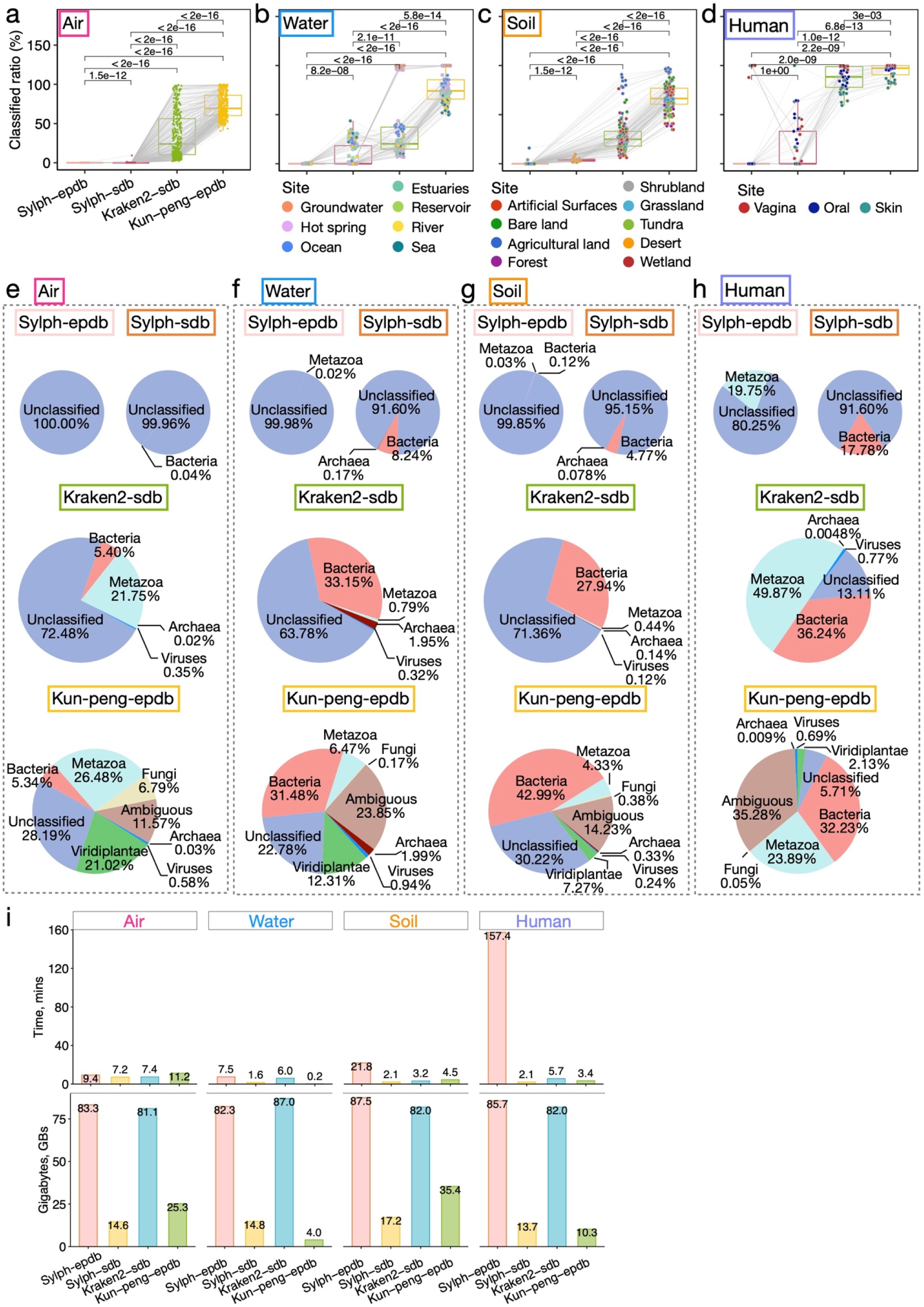
Comprehensive evaluation of Kun-peng against other classifiers on diverse environmental samples using standard and expansive pan-domain databases. a-d. Classification rate comparison across environmental samples from: (a) air (n=301), (b) water (n=105), (c) soil (n=135), and (d) human (n=45) sites. Tools were tested using different database configurations: expansive pan-domain database (Kun-peng-epdb, Sylph-epdb) and standard database (Sylph-sdb, Kraken2-sdb). P-values from pairwise comparisons using the Kruskal-Wallis test with post hoc Games-Howell tests are shown. e-h. Domain-level taxonomic composition analysis for representative samples from: (e) air, (f) water, (g) soil, and (h) human environments. Pie charts show the relative abundance of different taxonomic groups across different tools and database configurations. i. Computational performance comparison across sample types showing processing time (upper panel, minutes) and peak memory usage (lower panel, gigabytes) for each tool-database combination. Values represent averages per sample for each environmental category.

This significant performance deterioration from Sylph-sdb to Sylph-epdb reveals fundamental limitations in Sylph’s approach to expansive pan-domain classification. These limitations can be attributed to several critical aspects of Sylph’s algorithmic design: 1) its statistical algorithm and parameters were optimized primarily on CAMI2’s synthetic datasets with simple prokaryotic community structures, making it ill-suited for complex environmental samples; 2) its k-mer subsampling strategy, which retains only 1/200 of k-mers, inherently reduces sensitivity, particularly problematic for complex samples where many organisms are present at low abundance; and 3) its statistical model appears to break down when handling multi-domain reference databases. While Sylph demonstrates good performance for bacterial and archaeal profiling in well-characterized microbiomes like the human gut, our results indicate that it is fundamentally limited when applied to complex environmental samples requiring pan-domain classification. In contrast, Kun-peng’s database architecture and full k-mer indexing approach are better suited for comprehensive environmental metagenome classification using pan-domain references.

We further compared the computational performance among tested tools using ten cores with default settings (Fig. 3i). When using Kun-peng-epdb with the memory-efficient mode, the average classification times per sample were 11.2 minutes for air, 0.2 minutes for water, 4.5 minutes for soil, and 3.4 minutes for human samples. Notably, these processing times were only marginally increased compared to Kraken2 using its standard database (Kraken2-sdb, 81GB), despite the database in Kun-peng-epdb being 23 times larger in size. In contrast, Sylph-epdb showed slightly faster processing time for air samples (9.4 mins) but substantially slower processing time for water (7.5 minutes), soil (21.8 minutes), and human (157.4 minutes) samples. However, the comparison of processing speed for Sylph-epdb is largely inconsequential given its failure to classify most samples effectively.

In addition to processing speed advantages, the peak memory usage (Fig. 3i) of Kun-peng-epdb was efficient at 25.3 GB, 4.0 GB, 35.4 GB, and 10.3 GB for samples from air (n=301), water (n=105), soil (n=135), and human body sites (n=45), respectively. These numbers are substantially less than Sylph-epdb (83.3 GB, 82.3 GB, 87.5 GB, and 85.7 GB) and Kraken2-sdb (81.1 GB, 87.0 GB, 82.0 GB, and 82.0 GB). Notably, even the highest peak memory usage (35.4 GB) was still 473-fold less than Kraken2’s peak requirement for the expansive pan-domain database (1.85 TB). Kun-peng’s peak memory requirement is dictated by the size of database blocks (4GB) and intermediate data, which varies based on read length, number of reads, and classification rates. This remarkable reduction in memory usage while maintaining effective classification performance showcases Kun-peng’s exceptional efficiency in taxonomic classifications against large-scale databases. Last but not least, minimal or low memory requirements on most HPC platforms can greatly reduce the overall job queueing times, making the actual time savings even more substantial when classifying large amounts of samples.

In conclusion, Kun-peng demonstrates superior memory efficiency, processing speed, and classification accuracy compared to the mainstream read-based taxonomy classifiers such as Kraken2, Centrifuge, and Sylph, providing a universally accessible, fast, and accurate taxonomic classification solution for global researchers.

## Acknowledgments

We are grateful to our colleagues at the core facility of the Life Sciences Institute, especially the NECHO high-performance computing cluster. This research was partly supported by grants from the National Natural Science Foundation of China (NSFC) (82173645 and 82341109).

## Competing interests

The authors declare no competing interests.

## Methods

### Downloading Databases

Kun-peng incorporates an enhanced NCBI command-line tool for efficient genomic data retrieval (Fig. 1a). This utility supports:

1. Comprehensive database access: Downloads from RefSeq and GenBank across diverse taxonomic groups (archaea, bacteria, viruses, fungi, plants, humans, protozoa, vertebrate mammals, other vertebrates, and invertebrates).
2. Flexible assembly levels: Retrieval of complete genomes, chromosomes, scaffolds, and contigs.
3. Resumable downloads: Implements breakpoint resumption technology, allowing paused downloads to continue from the point of interruption.
4. Incremental updates: Enables downloading only new or updated files, conserving bandwidth and time.
5. Data integrity: Performs MD5 checksum verification for each file, ensuring uncorrupted transfers.

These enhancements significantly improved the database download functionality of previous tools, such as Kraken2, offering more robust and efficient data acquisition.

### Building the reference database

Kun-peng’s hash table for minimizer/LCA key-value pairs resembles Kraken2’s but divides the table into ordered 4GB blocks to reduce memory usage (Fig. 1a). The process generates a minimizer value vector using the k-mer method, sliding window algorithm, and hash function, then transforms it into cell records containing compact hash codes and LCA values. Unlike Kraken2, Kun-peng implements a uniqueness check during minimizer generation to retain only unique ℓ bp minimizers, preventing over-counting from identical sequence regions that could lead to false positives. The compact hash code’s bit count varies based on unique taxonomy ID numbers in the reference library.

Cell record position is determined by:

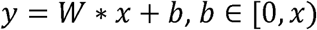

where *y* is the minimizer value, *x* is the full database capacity, *W* is the quotient of *y⁄x*, and *b* is the remainder and position of the cell record.

Kun-peng resolves storage conflicts by:

1. Using the ancestor taxonomy ID when y is identical but taxonomy IDs differ.
2. Applying linear probing (incrementing b by 1) when y differs but *b*1 = *b*2.

Kun-peng innovates by splitting the hash table into ordered 4GB blocks:

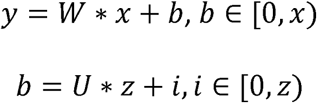

Where *U* is the block file count, *z* is the block capacity (4GB), and *i* determines the cell record’s position in a block.

The hash temporary file comprises block index (*U*), compact hash code, and LCA value, preserving all information and explaining consistent performance between Kun-peng-B (reference database divided into 4GB blocks) and Kun-peng-W (whole reference database without partitioning).

Our study with 42,485 genomes uses *x* = 56GB and *z* = 4GB, resulting in *U* = 14 blocks. Kun-peng loads one 4GB block at a time, storing others on disk, enabling large database handling on personal computers with low memory.

Notably, Kun-peng is compatible with Kraken2’s hash table, allowing direct database building by splitting into 4GB blocks.

### Classification of a sequence

Kun-peng’s sequence classification method builds upon Kraken2’s approach, incorporating key modifications to reduce false positives and implementing parallel processing algorithms to enhance processing speed. The process begins by generating a sample temporary file containing the minimizer vector index, block index, compact hash code, file number, and sequence number (Fig. 1b). For each k-mer in an input sequence, Kun-peng identifies its minimizer. If the minimizer matches a key in the compact hash code and block index, the associated LCA value is assigned to the k-mer. Kun-peng optimizes the sliding window algorithm by counting minimizers from the same sequence position only once, effectively reducing false positives. The classification then proceeds similarly to Kraken2, mapping k-mer hits to taxa, constructing a pruned classification tree, and classifying the sequence using the leaf of the maximally scoring root-to-leaf path.

To accommodate diverse computing environments, Kun-peng offers two operational modes:

1. Memory-Efficient Mode (Kun-peng-M): This mode processes one block of the reference database at a time, requiring memory slightly above 4GB (block size). It is particularly suitable for environments with limited memory resources.
2. Full-Speed Mode (Kun-peng-F): This mode loads all database blocks simultaneously using multithreading, requiring memory slightly larger than the entire hash table. While Kun-peng-F offers faster processing, it is more suitable for high-memory environments.

These dual modes provide flexibility, allowing Kun-peng to operate efficiently across a wide spectrum of computational setups, from personal computers to high-performance computing platforms. This adaptability ensures that researchers can leverage Kun-peng’s capabilities regardless of their available computational resources.

### Comparison of benchmark performance and computational resource usage

We evaluated Kraken2 (v2.1.3), Centrifuge (v1.0.4), and Sylph (v0.6.1) using two mock communities with 14 samples: Amos HiLo and Mixed mock communities^21^ and NIST mock communities^22^. The expected relative abundances are listed in Tables S1 and S2, with sequence data accessible through NCBI SRA (accession numbers SRR11487931-SRR11487935 for HiLo, SRR11487937-SRR11487941 for Mixed, and study accession SRP436666 for NIST).

We constructed a standard database for each tool from the complete RefSeq genome (January 1, 2020), covering 904 archaeal, 68,743 bacterial, and 14,033 viral genomes. All databases were built using each tool’s default parameters. For metagenomic classification, Kraken2 and Kun-peng were executed with a confidence score of 0.3, while Centrifuge was run with default settings. The classifications from Kraken2, Kun-peng, and Centrifuge were processed with Bracken (v2.9) for abundance estimation, and taxa with abundances below 0.01% were filtered out to minimize noise ^8^. Sylph was run with its default settings.

Precision was calculated as the ratio of correctly identified genera or species to the total number of taxa identified by each tool (true positives divided by true positives plus false positives). Sensitivity was calculated as the ratio of correctly identified genera or species to the total number of true genera or species present in the sample (true positives divided by true positives plus false negatives). We generated the area under the precision-recall curve (AUPR) by varying the abundance threshold from 0 to 1.0 ^8^.

Time and memory consumption were profiled on a Zhejiang University HPCP using 10 threads. We used the BMocK12 dataset^23,25^ (NCBI SRA accession SRR8073716), subsampled to six 25- million read sets using seqtk (https://github.com/lh3/seqtk). The seqtk sampling command was executed for both forward and reverse reads, varying the seed from 100 to 600 in increments of 100. Additionally, we profiled Kun-peng-M on a MacBook Pro (32GB memory, Intel Core i5) and a Macstudio (32GB memory, Apple M1 Max), with or without a Samsung 990 PRO SSD, estimating time usage for both environments.

We constructed an expansive pan-domain database from GenBank and RefSeq complete genomes and chromosomes (as of August 6, 2024), encompassing archaea (1,411 genomes), bacteria (105,560), fungi (1,705), human (2), invertebrate (226), plant (2,153), protozoa (43), vertebrate mammalian (128), vertebrate other (286), and viral (193,810) genomes. This extensive database (4.3TB of fna sequences) generated a 1.85TB hash index for Kun-peng and an 81GB index for Sylph. For comparison, we built a standard database using RefSeq data from the same data, containing only archaea (619), bacteria (46,433), human (2), and viral (14,972) genomes, which produced an 81GB hash index for Kraken2. We also evaluated Sylph using its pre-built standard database GTDB-R220 in 14GB, which contains 113,104 bacterial/archaeal species representative genomes. To evaluate classification performance, we analyzed 586 samples, including 301 air exposure samples ^26^, 105 water samples from estuaries ^27^, groundwater ^28^, hot spring ^29^, ocean ^30^, reservoir ^31^, rivers ^32^, and sea ^32^, 135 soil samples ^32,33^ from agricultural land, artificial surfaces, bare land, forest, grassland, shrubland, tundra, wetland, and desert, and 45 samples from human body site, including oral ^34^, skin ^35^, and vagina^36^. Except for air exposure samples, each site contributed 15 samples, with download sources detailed in Table S3. We processed all samples using four configurations: Kun-peng with the pan-domain database (Kun-peng-epdb), Sylph with both pan-domain and standard databases (Sylph-epdb and Sylph-sdb), and Kraken2 with the standard database (Kraken2-sdb). All tools were run with default parameters, except for Sylph which was run with the-u option to obtain true sequence abundances. This experimental design enabled evaluation of how comprehensive pan-domain databases impact classification performance across diverse environmental samples with different tools.

Kun-peng’s peak memory requirement is determined by two main factors: the size of database blocks (4GB) and the intermediate data. The size of intermediate data is primarily influenced by read length, number of reads, and classification rates. Longer reads, a higher number of reads, and higher classification rates all contribute to larger intermediate data, potentially increasing memory usage. To optimize memory efficiency, we employed Kun-peng-M with a batch size of 16 when classifying the exposure samples, which effectively divides the large intermediate data into 16 manageable blocks. This strategy further significantly reduces memory consumption.

## Notes

https://github.com/eric9n/Kun-peng

### Competing Interest Statement

The authors have declared no competing interest.

