## Supplemental information for "Kun-peng: an ultra-memory-efficient, fast, and accurate pan-domain taxonomic classifier for all"

^4^Lee Kong Chian School of Medicine, Nanyang Technological University, Singapore. School of Chemistry, Chemical Engineering and Biotechnology, Nanyang Technological University, Singapore.

^#^Contributed equally

**Table S1**. Expected relative abundance for the Amos HiLo and Mixed mock communities.

| Species | Genera | Phyla | Taxid | HiLo,  RA, % | Mixed,  RA, % |
| --- | --- | --- | --- | --- | --- |
| Akkermansia muciniphila | Akkermansia | Verrucomicrobiota | 239935 | 0.18 | 6.37 |
| Alistipes finegoldii | Alistipes | Bacteroidota | 214856 | 1.30 | 4.54 |
| Anaerostipes hadrus | Anaerostipes | Bacillota | 649756 | 1.75 | 6.11 |
| Bacteroides thetaiotaomicron | Bacteroides | Bacteroidota | 818 | 7.72 | 2.69 |
| Bacteroides uniformis | Bacteroides | Bacteroidota | 820 | 1.05 | 3.66 |
| Bifidobacterium longum | Bifidobacterium | Actinomycetota | 216816 | 37.02 | 12.92 |
| Blautia wexlerae | Blautia | Bacillota | 418240 | 0.11 | 3.77 |
| Clostridium butyricum | Clostridium | Bacillota | 1492 | 10.59 | 3.70 |
| Collinsella aerofaciens | Collinsella | Actinomycetota | 74426 | 1.99 | 6.95 |
| Escherichia coli | Escherichia | Pseudomonadota | 562 | 9.33 | 3.26 |
| Eubacterium hallii | Eubacterium | Bacillota | 39488 | 1.48 | 5.16 |
| Faecalibacterium prausnitzii | Faecalibacterium | Bacillota | 853 | 0.16 | 5.49 |
| Lactobacillus gasseri | Lactobacillus | Bacillota | 1596 | 0.26 | 8.97 |
| Parabacteroides distasonis | Parabacteroides | Bacteroidota | 823 | 10.10 | 3.52 |
| Prevotella copri | Prevotella | Bacteroidota | 165179 | 13.84 | 4.83 |
| Prevotella melaninogenica | Prevotella | Bacteroidota | 28132 | 1.53 | 5.34 |
| Roseburia hominis | Roseburia | Bacillota | 301301 | 1.35 | 4.72 |
| Roseburia intestinalis | Roseburia | Bacillota | 166486 | 0.11 | 3.88 |
| Ruminococcus gauvreauii | Ruminococcus | Bacillota | 438033 | 0.12 | 4.13 |

**Table S2.** Expected relative abundance for the NIST communities.

| Pool | Species | genera | Phyla | Taxid | RA, % |
| --- | --- | --- | --- | --- | --- |
| Pool A | Escherichia coli | Escherichia | Pseudomonadota | 83334 | 0.51 |
|  | Staphylococcus aureus | Staphylococcus | Bacillota | 1280 | 0.28 |
|  | Neisseria meningitidis | Neisseria | Pseudomonadota | 487 | 0.21 |
| Pool B | Salmonella enterica | Salmonella | Pseudomonadota | 28901 | 0.33 |
|  | Acinetobacter baumannii | Acinetobacter | Pseudomonadota | 470 | 0.26 |
|  | Klebsiella pneumoniae | Klebsiella | Pseudomonadota | 573 | 0.41 |
| Pool C | Streptococcus pyogenes | Streptococcus | Bacillota | 1314 | 0.16 |
|  | Achromobacter xylosoxidans | Achromobacter | Pseudomonadota | 85698 | 0.48 |
|  | Shigella sonnei | Shigella | Pseudomonadota | 624 | 0.36 |
| Pool D | Enterococcus faecalis | Enterococcus | Bacillota | 1351 | 0.30 |
|  | Vibrio furnissii | Vibrio | Pseudomonadota | 29494 | 0.45 |
|  | Listeria monocytogenes | Listeria | Bacillota | 1639 | 0.25 |

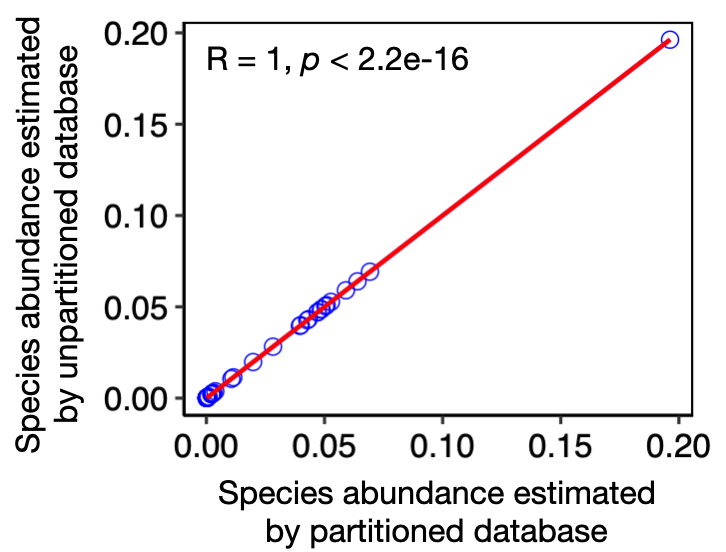

**Fig. S1.** Correlation of classified species abundance between partitioned and unpartitioned databases in Kun-peng for two mock datasets. The correlation coefficient (R) and p-value were calculated using Spearman's rank correlation.
